# Populating a Continent: Phylogenomics Reveal the Timing of Australian Frog Diversification

**DOI:** 10.1101/2023.05.03.539251

**Authors:** Ian G. Brennan, Alan R. Lemmon, Emily Moriarty Lemmon, Conrad J. Hoskin, Stephen C. Donnellan, J. Scott Keogh

## Abstract

The Australian continent’s size and isolation make it an ideal place for studying the accumulation and evolution of biodiversity. Long separated from the ancient supercontinent Gondwana, most of Australia’s plants and animals are unique and endemic, including the continent’s frogs. Australian frogs comprise a remarkable ecological and morphological diversity categorized into a small number of distantly related radiations. We present a phylogenomic hypothesis based on an exon-capture dataset that spans the main clades of Australian myobatrachoid, pelodryadid hyloid, and microhylid frogs. Our time-calibrated phylogenomic-scale phylogeny identifies great disparity in the relative ages of these groups which vary from Gondwanan relics to recent immigrants from Asia and include arguably the continent’s oldest living vertebrate radiation. This age stratification provides insight into the colonization of, and diversification on, the Australian continent through deep time, during periods of dramatic climatic and community changes. Contemporary Australian frog diversity highlights the adaptive capacity of anurans, particularly in response to heat and aridity, and explains why they are one of the continent’s most visible faunas.

## Introduction

Frogs are an ancient vertebrate radiation originating in the Permian more than 250 million years ago (Hime et al. 2021). They share a unique and unusual morphology yet are a spectacularly successful group, with more than 7,500 extant species spread across most of the world (AmphibiaWeb 2022). Despite their age, much of this diversity, potentially more than 95%, has developed since the Cretaceous-Paleogene mass extinction (65 mya) (Feng et al. 2017). Australia is one of the driest continents on Earth yet, surprisingly, it is home to nearly 250 frog species. Australia’s frogs belong to just four anuran groups spread widely across the “modern frog” suborder Neobatrachia: (1) Myobatrachoidea comprising the Limnodynastidae (66 species) and Myobatrachidae (70 spp.); (2) Hyloidea represented by the family Pelodryadidae (91 spp.); (3) the Microhylidae subfamily Asterophryinae (24 spp.); and (4) a single Ranidae species in the genus *Papurana*. These groups show very different levels of species richness and geographic spread across the continent (Fig.1). However, together they have radiated to inhabit almost every part of Australia including tropical rainforests, alpine streams, featureless boulder piles, and hyper-arid deserts.

**Figure 1.**
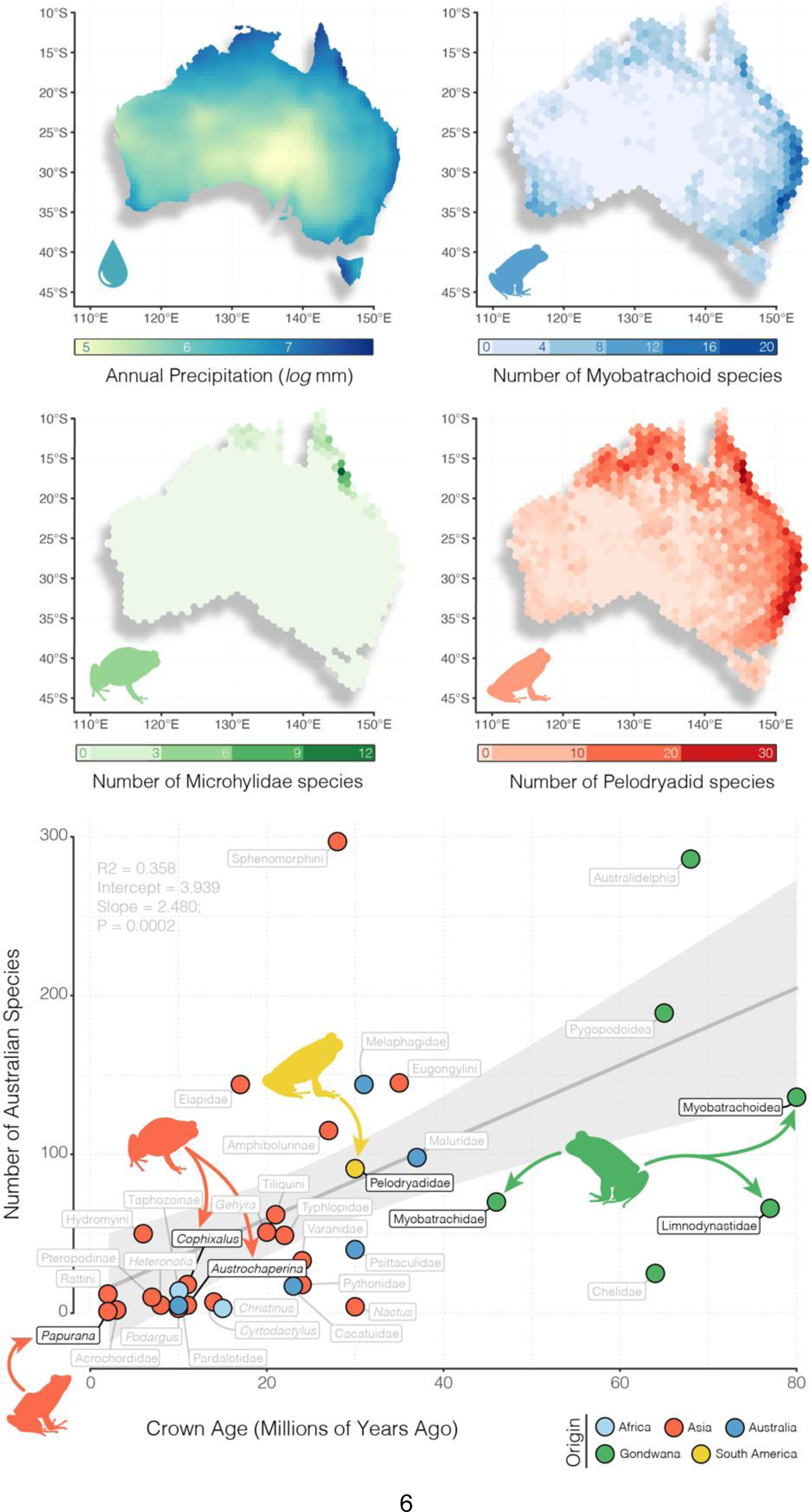
Australian frogs show an imbalance in species richness, age, and geographic spread. Above, maps of richness for the three focal radiations (with Limnodynastidae and Myobatrachidae presented together as Myobatrachoidea) represent visually how contemporary patterns of frog richness reflect water availability, and are highest in the wet temperate, subtropical, and tropical rainforests of the east coast. We show annual precipitation here for ease of interpretation but Australian frog richness is potentially better explained by actual evapotranspiration (Coops et al. 2018). Species occurrence records were collated from the Atlas of Living Australia (https://ala.org.au). Below, Australian radiations can be divided broadly into (1) relictual Gondwanan clades >40 myo (green), (2) ancient colonizing groups (>20 myo, <40 myo; varied colors), or (3) immigrant clades of Asian origin (orange). Each point is colored according to the region of hypothesized origin and labeled by the narrowest phylogenetic taxonomy. Black labels indicate focal groups and grey labels indicate other Australian vertebrate clades. Regression in background is fit to all points with the exception of Limnodynastidae and Myobatrachidae (included jointly as Myobatrachoidea) and shows a general pattern of increasing species richness with age. This pattern holds equally for a regression of just frog clades (R^2^=0.849, intercept=1.827, slope=1.805, p=0.016).

While we know a great deal about many aspects of Australian frog biology (Tyler 1998; Anstis 2017), the age of each of the major groups and the timing of their subsequent diversification, is poorly understood. Since the origin of frogs over 250 million years ago, the landmass that is now Australia has traveled extensively. Long ago it was part of the supercontinent Pangea before separating as a component of Gondwana alongside South America, Africa, Antarctica, and India. Sometime around 50 million years ago Australia separated from Antarctica and began drifting alone towards Asia (Hall 2002; Bijl et al. 2013). Given the long evolutionary history of frogs, and Australia’s varied geographic affinities with other landmasses, we ask three related questions: (1) Where did Australia’s frogs originate? (2) When did they get to Australia? and (3) Who and where are their closest relatives? Answering these questions provides context for the varied species richness and ecological diversity of these groups and offers important insight into the evolution of a continental fauna.

## Materials and Methods

We assembled an exon-capture dataset comprising 99 frog species spanning all major anuran clades and with particular focus on the families Pelodryadidae, Microhylidae, Limnodynastidae and Myobatrachidae (Table S1). This dataset includes near-complete (92%) genus-level sampling of Australia’s frogs. We generated new Anchored Hybrid Enrichment (AHE—Lemmon et al. 2012) data for 83 samples and combined these with outgroup samples from Hime et al.’s (2021) amphibian phylogenomic dataset. Outgroup sampling was designed around maximizing commonly used anuran fossil calibrations to provide a consistent time-calibrated phylogenomic estimate of Australian frogs. Data from different AHE projects were combined using custom scripts which relied on *metablastr* to identify orthologous loci (*blast_best_reciprocal_hit*) (Benoit & Drost 2021), *mafft* to align them (*--add*, *--keeplength*) (Katoh et al. 2013), and *AMAS* to manipulate alignments (Borowiec 2016). We reconstructed individual genealogies for our exon-capture data (n = 450) under maximum-likelihood in IQTREE (Nguyen et al. 2015), allowing the program to assign the best fitting model of nucleotide substitution using ModelFinder (Kalyaanamoorthy et al. 2017) and then perform 1,000 ultrafast bootstraps (Minh et al. 2013). We then estimated a species tree using the quartet-based summary method ASTRAL III (Zhang et al. 2018) with IQTREE gene trees as input. To complement our coalescent-consistent summary method we also estimated a species tree from the concatenated alignment using the edge-unlinked partition model GHOST implemented in IQTREE. This allowed us to more accurately model rate variation among sites and samples. To estimate divergence times among taxa on the ASTRAL species tree we applied a series of fossil calibrations first compiled by Feng et al. (2019) (Table S2) and used the Bayesian divergence time software MCMCtree (Rannala & Young 2007). We started by concatenating all exonic loci (n=390; Supp. *Sequence Identity*) and partitioning them into two partitions, first and second codons together, and third codons separately, following the strategy of dos Reis et al. (2018). Complex partitioning strategies such as filtering by evolutionary rate are possible but less influential than the absolute number of partitions (dos Reis et al. 2012). Additional data partitions ultimately incur substantial computational costs for modest increases in dating precision, and so we opted instead for a more conservative approach. We then used *baseml* to estimate approximate likelihoods (dos Reis & Yang 2011) and branch lengths before running *mcmctree* on the gradient and Hessian (in.BV file) for ten replicate analyses. We inspected mcmc files for stationarity and compared for convergence, then combined them using logCombiner, and used this combined mcmc file to summarize divergence times on our tree (*print = −1* in .ctl file). Sample, alignment, and gene tree summary statistics are presented in Supplementary Material (Fig.S1-3) and are available alongside all other materials on Dryad (doi:10.5061/dryad.zpc866tcj) and GitHub (https://github.com/IanGBrennan/Crown_Frogs).

To investigate the biogeographic origins of Australian frogs we reconstructed ancestral ranges using *BioGeoBEARS* (Matzke 2014). The deep timescale of frog evolutionary history necessitates accounting for continental rearrangement and dispersal barriers by incorporating time-stratified information from plate tectonics. To accomplish this we designed a series of models that augment dispersal probability as a function of distance among areas and adjacency. Briefly, these models penalize dispersal probability as distance between areas increases, and as the *type* of distance changes (e.g. over-land vs. over-water dispersal). To identify the dispersal path of the pelodryadid tree frogs and how they arrived in Australia from a South American ancestor (Pyron 2014), we designed two data sets. The first requires the Pelodryadidae to have travelled from South America through Antarctica and into Australia (*H1*) and the second allows an overwater dispersal directly from South America to Australia (*H2*). Comparative model fit was assessed via AIC. Model specifics can be found in the *Supplementary Materials and Methods*.

## Results

Species tree topologies are nearly identical across the quartet-based coalescent method (ASTRAL) and concatenation under the GHOST heterotachy model (IQTREE), and are broadly consistent with previously published phylogenomic frog hypotheses (Feng et al. 2017; Streicher et al 2018; Streicher et al. 2020; Hime et al. 2021) (Fig.2, S4—S6). We estimate well-supported phylogenies with few unresolved nodes among Australian taxa. Australian microhylids fall into two non-sister clades, each nested within the primarily New Guinean Asterophryinae. Pelodryadids have diverged into two to three deep groups, with *Cyclorana* and *Nyctimystes* embedded within divergent clades of *Litoria*. Ancient splits among myobatrachoids show some uncertainty with a paraphyletic estimation of the Myobatrachidae. There is strong support uniting the genera *Mixophyes* and *Rheobatrachus*, and moderate support (LPP 90) places this myobatrachid clade as sister to the Limnodynastidae, to the exclusion of remaining myobatrachid genera.

**Figure 2.**
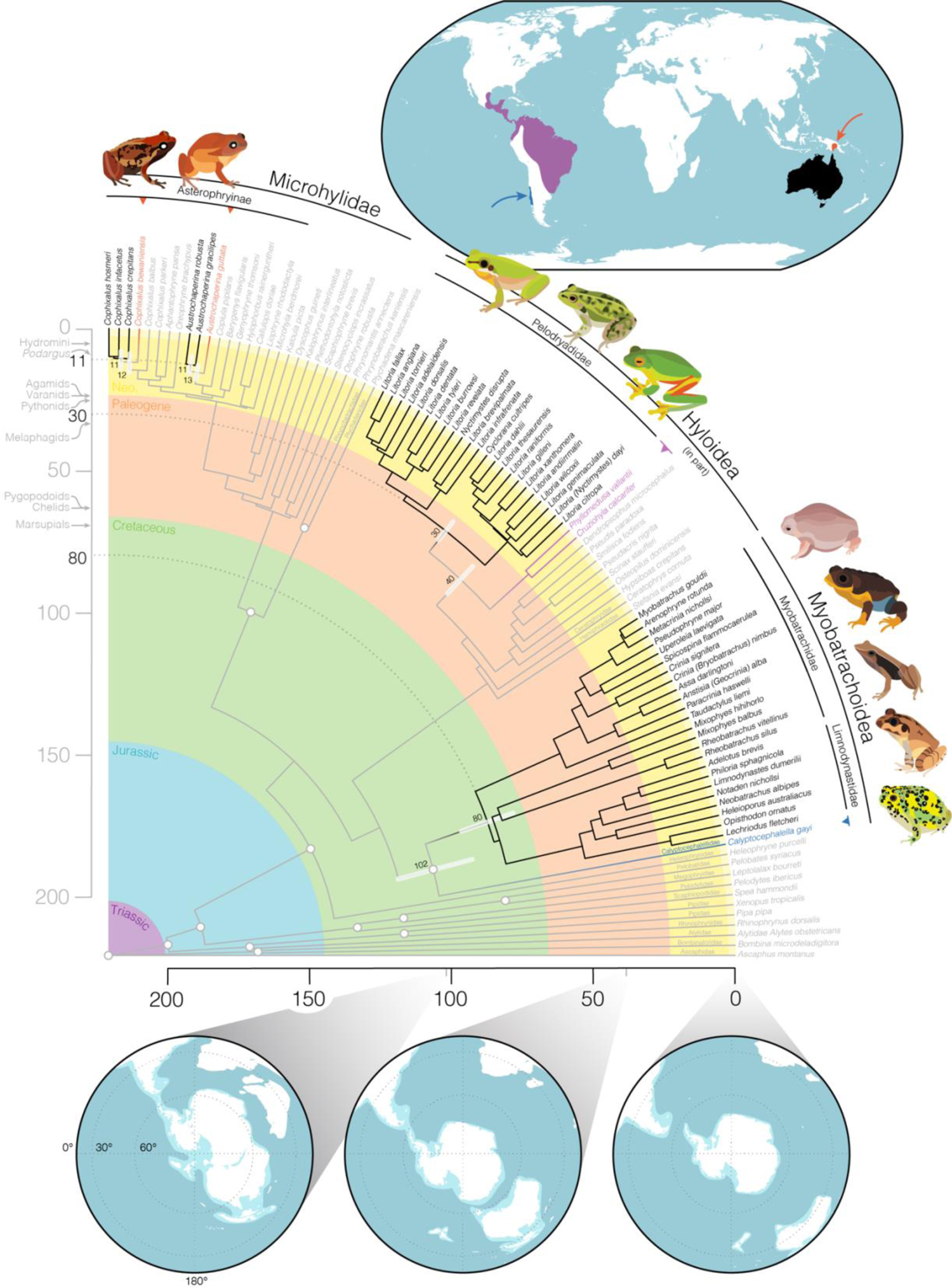
Time-calibrated frog phylogeny highlights the varied origins and staggered arrival of the four major frog families that comprise the Australian anuran fauna. Primarily Australian clades are identified by black branches and text, their closest living relatives outside of Australia are noted by colored branches and text, and outgroup taxa are grey. White circles at nodes identify the location of fossil calibrations (see Table S2). Upper inset map shows the general geographic location of: (red) closely related microhylids in New Guinea, (purple) phyllomedusid hylids in South America, and (dark blue) *Calyptocephallela* in Chile. Lower inset maps show the connection and proximity of Australia to other Gondwanan continents as Australia drifted away over the past 100 million years. White indicates contemporary coastlines, light blue the continental plates, and dark blue the oceans. Maps were generated using GPlates and input files modified from Landis (2017). Partial fan phylogeny was plotted using *phytools* in the R programming environment. Annotations on vertical time axis show the age of crown divergences of other notable Australian groups for temporal context (see Fig.1). Species illustrated clockwise from top left: *Cophixalus infacetus, Austrochaperina robusta, Litoria fallax, Litoria dahlii, Litoria xanthomera, Myobatrachus gouldii, Spicospina flammocaerulea, Taudactylus acutirostris, Mixophyes balbus, Notaden bennettii*.

Concatenated and coalescent topologies differ at three very short branches which bear no significant implications for our understanding of the relationships of Australian frogs (Fig.S5). Successive short branching events such as these are known to mislead tree inference from concatenated data, and so are not surprising (Linkem et al. 2016). We find support in the GHOST model for four distinct rate classes, which vary in total tree length (TTL) by more than 50x, providing evidence of strong heterotachy among sites. The distribution of TTL among branches across the four trees however, is largely consistent suggesting little effect of heterotachy among lineages.

Crown divergences of the three Australian frog radiations can be clearly separated into old (Myobatrachidae and Limnodynastidae–80 mya), intermediate (Pelodryadidae–30 mya), and young (Asterophryinae–11 mya) (Fig.2). The youngest Australian group, microhylids in the genera *Austrochaperina* and *Cophixalus*, are embedded deeply within the subfamily Asterophryinae and appear to represent two separate, relatively recent (≃11 mya) dispersals into Australia from New Guinea. Pelodryadidae tree frogs also share a complex biogeographic history across Australasia, with several species groups split across the Torres Strait (separating Australia and New Guinea), suggesting frequent biotic exchange. However, the origins of the Pelodryadidae are far older. Their closest extant relatives are the iconic Phyllomedusidae found throughout Central and South America, with the crown split between extant Pelodryadidae in Australia/New Guinea and South America estimated at approximately 40 million years ago. Australian myobatrachids and limnodynastids also have their closest living relatives in South America—the Calyptocephallelidae, represented here by *Calyptocephallela*, the Helmeted Water Toad of Chile. The crown split between extant myobatrachoids in Australia and calyptocephalellids in Chile is ancient, occurring more than 100 million years ago.

Biogeographic modelling provides support for a diversification scenario in which the dispersal of frogs was influenced by vicariant events (parameter *j*), distance among biogeographic regions (*x*), and dispersal type (*w*; over-land vs. over-water) (Table S3). The top two models account for more than 80% of AIC weight, and both correspond to pelodryadid dispersal Hypothesis 1 in which treefrogs dispersed through Antarctica to reach Australia (DEC+j+x+w *H1*, AICw 59.7; DEC+j+x *H1*, AICw 21.5). The preferred model represents a meaningful improvement over similar models under a pelodryadid dispersal Hypothesis 2 (Fig.3, S7; Table S3). Parameter estimates of *x* under the top two models suggest that doubling the distance between areas reduces dispersal probability by one-third to one-half. Parameter estimation of *w* under the preferred model suggests that overland dispersal probability among non-adjacent areas is one-third that of between adjacent areas, and overwater dispersal probability is just one-tenth.

**Figure 3.**
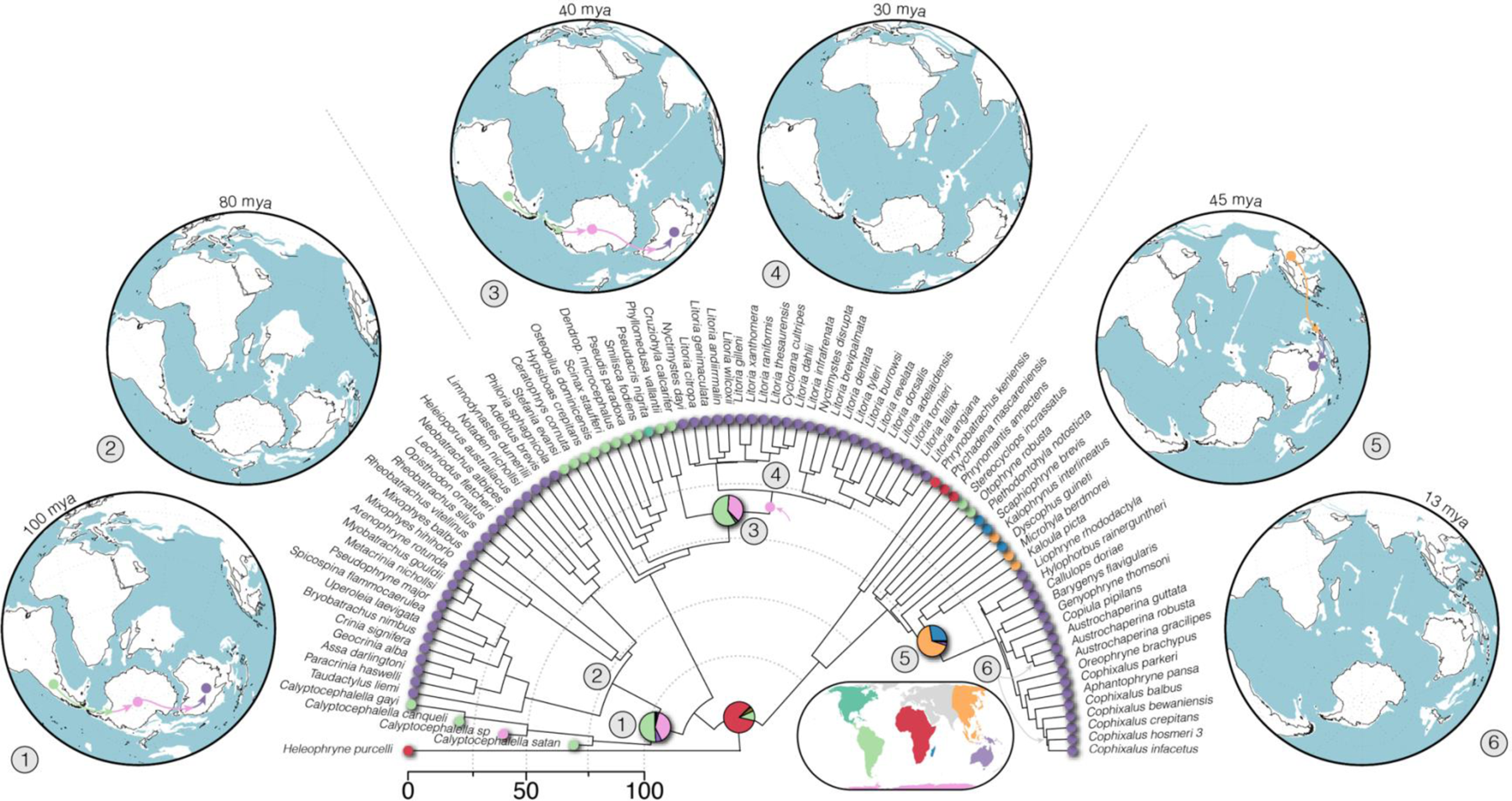
Simplified biogeographic history of Australian frogs with a focus on the range reconstruction of their immediate ancestors (complete figure in Fig.S7). Ranges have been estimated under the preferred model DEC+*j*+*x*+*w* supporting Hypothesis 1 (Antarctic dispersal of Pelodryadidae frogs; pink arrow on tree indicates ancestral pelodryadid constrained to Antarctica) in *BioGeoBEARS*. Pie charts represent range probability at nodes with colors corresponding to inset map. Circular world maps show geological reconstructions at relevant time points, with numbers mapped to nodes of interest. Colored arrows indicate hypothesized dispersal paths for each clade. Under this biogeographic model the ancestors of both the Myobatrachoidea and Pelodryadidae lived in South America, and Australo-Papuan microhylids (Asterophryinae) originate from an Asian ancestor. The most likely dispersal path for the Pelodryadidae included expansion across Antarctica after divergence from the Phyllomedusidae. Phylogeny plotted with *phytools*, maps generated by the Ocean Drilling Stratigraphic Network (https://www.odsn.de/odsn/services/paleomap/paleomap.html).

Ancestral range reconstructions provide evidence that both myobatrachoid and pelodryadid frogs are descended from South American ancestors. Asterophryinae microhylids, in which the Australian microhylids are embedded, likely diverged from an ancestor found in Asia.

## Discussion

Here we present the first reliable estimates of relationships among nearly all of Australia’s native frog genera (25 of 27) and major clades of the diverse genus *Litoria*. Our investigation into the timing and origins of the Australian frog fauna reveals a staggered colonization and population of the continent. This stratified arrival and radiation of Australian frogs took place under the varied environmental conditions of vastly different eras. Across these eras Australia has flourished through a warm and wet Eocene, cooling and drying following the onset of Antarctic glaciation in the Oligocene, warm and forested Miocene, and a gradual aridification leading to its present status (Byrne et al. 2011, Pross et al. 2012, Macphail & Hill 2018, Mao & Retallack 2019).

### Origins and Biogeography

The Myobatrachidae and Limnodynastidae (together—myobatrachoids) represent the oldest, most diverse (136 spp.), and only near-endemic of Australia’s frog radiations (4 spp. are found in New Guinea). They share a long history with South America and its Gondwanan past. Anchored by a deep split with the South American *Calyptocephalella* (roughly 100 mya; Fig.2), early divergences among the myobatrachoids, principally between *Mixophyes*, *Rheobatrachus*, and the limnodynastids, occurred in the late Cretaceous (80–70 mya), preceding the isolation of Australia from Antarctica. This dates to a time when South America, Antarctica, and Australia were a continuous landmass that was likely temperate in climate (Palazzesi & Barreda 2007; Mörs et al. 2020). The phylogenetic depth and distribution of myobatrachoids and calyptocephalellids across these now widely disjunct continents suggests a historically continuous distribution across southern Gondwana, including Antarctica. This idea is supported by the recent discovery of an extinct calyptocephallelid from mid-Miocene Antarctica that lived more than 40 mya (Mörs et al. 2020). The persistence of calyptocephalellids in Antarctica into the Late Eocene highlights the dichotomy between young extant myobatrachid and limnodynastid diversity (most species < 30 mya) and ancient splits between limnodynastids and myobatrachids and within myobatrachids (> 70 mya). The tips of these long branches are likely the survivors of a much greater southern Gondwanan myobatrachoid diversity, potentially mirroring the diversity of extinct calyptocephalellids through southern South America and Patagonia (Nicoli et al. 2022).

Australian myobatrachoids however are not the only group with close connections to South America. The Pelodryadidae are a species rich (>220 spp.) and morphologically diverse clade of Australasian frogs. Embedded within the primarily Neotropical treefrogs, they show a more recent late-Eocene divergence from their South American relatives the Phyllomedusidae, some 40 mya. Crown divergence of the pelodryadids occurred in the mid-to-late Oligocene (30 mya) before erupting into a radiation across Australia and New Guinea in the early Miocene. This timing has spurred speculation about the origins of pelodryadids either as part of a young Gondwanan group or more recent over-water dispersers from South America (Pyron 2014). Divergence between phyllomedusids and pelodryadids 40 mya aligns with the opening of the Drake Passage and separation of South America from Antarctica (Toumoulin 2020). Unfortunately, this does not provide any certainty about how pelodryadids arrived in Australia. While the Brazil Current would have provided a favorable trajectory for rafting frogs, the over-water distance between South America and Australia remained immense. Our biogeographic modelling indicates that the probability of overwater dispersal is just a fraction of that overland, making rafting seem improbable. Instead, we suggest a more likely scenario is that pelodryadids dispersed from South America through Antarctica and into Australia (Fig.3). Climate reconstructions suggest warm temperate/tropical habitats across Antarctica which would have been suitable through a long period of the Eocene (Pross et al. 2012). Dispersal via Antarctic land bridges would have had to occur prior to the Eocene-Oligocene cooling (34 mya) that blanketed Antarctica beneath an ice sheet (van den Ende et al. 2017).

Contrasting with the comparatively ancient limnodynastids, myobatrachids, and pelodryadids, Australia’s youngest anuran radiation are the microhylids. Embedded deeply in the Asterophryinae subfamily, two similarly aged clades (12–13 mya) of *Austrochaperina* and *Cophixalus* crossed the gap from New Guinea to Australia in the mid Miocene. This time frame coincides with a period of increased variation in sea surface levels driven by cooling global temperatures following the mid Miocene climatic optimum. Dropping sea levels likely repeatedly exposed a landbridge between southern New Guinea and northern Australia (both Cape York and the Top End) and facilitated biotic exchange between these landmasses (Mitchell et al. 2014). The young age of these clades, and existence of two other species-rich incumbent frog clades in the pelodryadids and myobatrachoids potentially explains why Australian microhylids are relatively species poor (*Austrochaperina*—5 spp., *Cophixalus*—18 spp.) and morphologically conservative compared to their New Guinean neighbors (200+ spp.), reflecting a pattern seen in monitor lizards (Pavón-Vázquez et al. 2021).

The sole Australian ranid *Papurana daemeli* is native but not endemic to the continent, and can be found broadly across Australo-Papua, extending to just beyond the edge of the Sahul shelf (Reilly et al. 2022). It belongs to a clade of frogs distributed throughout southeast Asia, Wallacea, and Sahul, with other *Papurana* species found in New Guinea and the Solomon Islands (Oliver et al. 2015; Chan et al. 2020). Though not included in our phylogenomic sampling, *Papurana daemeli* is likely a relatively young species (<7 mya) with limited divergence between populations found in Wallacea and Sahul (Reilly et al. 2022). The broad distribution of *P. daemeli* across Australo-Papua suggests either a very recent colonization of Australia or vicariant speciation followed by subsequent dispersal out of Australia and back into New Guinea and Wallacea.

The staggered temporal origins of Australian frogs exemplifies the general colonization history of Australian vertebrates. Radiations of mammals, birds, frogs, and reptiles fall into discretized temporal groups broadly identified as (1) Gondwanan relics >40 myo, (2) old established clades (20—40 myo) with varied origins, or (3) recent immigrants from Asia (<20 myo). The Limnodynastidae and Myobatrachidae fall undoubtedly into the Gondwanan group alongside ancient Australian radiations like Australidelphian marsupial mammals which include koalas, kangaroos, and Tasmanian devils; side-necked chelid turtles; and pygopodoid geckos which include the bizarre limbless pygopodids. These groups—with the exception of pygopodoids—have close links to South American relatives based on molecular and fossil evidence (Georges et al. 1999; Mitchell et al. 2014). While a Pelodryadidae link with South America is clear, they are perhaps the sole radiation to have emigrated from South America to Australia since the continental breakup. Most other similarly aged Australian groups instead show signal of Asian or Australian origins. In comparison, the Australian microhylids (*Austrochaperina*, *Cophixalus*) and the ranid *Papurana daemeli* are relatively young arrivals from New Guinea with deeper origins in Asian groups. Both the Asterophryinae and Ranidae, to which these species belong, have a long history in the Sunda and Wallacean regions, reflecting patterns of old diversity in this tectonically active area. Alongside a number of other groups such as pythons (Esquerré et al. 2020), monitor lizards (Brennan et al. 2021), honeyeater birds (Marki et al. 2017), dragon lizards (Tallowin et al. 2020), elapid snakes (Keogh 1998), various gekkonid gecko genera (Heinicke et al. 2011), megabats (Tsang et al. 2020), frogmouth birds (Oliver et al. 2020), cockatoos and parrots (Schweizer et al. 2011), several skink subfamilies (Skinner et al. 2011), and two rodent groups (Roycroft et al. 2020), they share diversity across Australia and New Guinea with repeated exchange between the two islands. Many of these groups show a telltale stepping stone biogeographic pattern that links them back to mainland Asian ancestors, with Australo-Papuan members deeply phylogenetically nested. In general, these Australian clades show a pattern of increasing species richness with clade age, however the drivers of such a pattern are potentially idiosyncratic (Fig.1) (Wiens 2011; Rabosky et al. 2012).

### Macroevolutionary Patterns

The radiation of frogs in Australia has occurred over a deep timescale and across a changing climatic landscape. Old species-poor lineages have become confined to the mesic-temperate fringes of the continent, while new niches and species have popped up in the expanding arid zone (Morgan et al. 2007; Novikova et al. 2020). And while frogs are found across most of the Australian continent, their basic moisture requirements and desiccation sensitivity mean that Australian amphibian diversity shows a stark mesic-arid gradient (Fig.1), similar to that seen for birds and mammals, and the inverse of lizards (Powney et al. 2010; Coops et al. 2018). Not all has been lost in the arid center though—several independent clades of dry-country inhabitants have evolved among Australia’s harsh sandy and stony deserts. *Neobatrachus*, *Notaden,* and *Cyclorana* have all evolved to aestivate through the hottest and driest seasons. These genera (commonly known as the water-holding frogs) are capable of growing epidermal cocoons to retain moisture that may see them through periods of extreme drought lasting from months to years (van Beurden 1980).

Along with changes in habitat and ecology, Australia’s frogs have also accumulated vast diversity in reproductive strategy, ontogenetic trajectory, and morphology (Crump 2015, Duellman 1992, Sherratt et al. 2018). While we do not present data on these topics, our well-resolved phylogenetic hypothesis provides new context for the macroevolution of some of these extreme traits. Unique rearing habits such as raising young in stomachs (*Rheobatrachus*), hip-pockets (*Assa*), or subterranean nests (*Myobatrachus*) exist on both long branches and deeply nested taxa suggesting a remarkable frequency of transition among states. Similarly, morphological variation has rapidly evolved to dramatic extremes. The long limbed highly aquatic *Litoria dahlii* with webbed feet and dorsally situated eyes is sister to the short-limbed burrowing water-holding frogs *Cyclorana* (Vidal-Garcia & Keogh 2015). Together these frogs are embedded deeply within the otherwise toe-padded and arboreal tree frogs, highlighting the adaptive capacity of pelodryadids. Myobatrachoids too have taken ecomorphology to the extreme, offering us what is perhaps the world’s strangest living anuran, the turtle frog *Myobatrachus gouldii*. In pursuit of their backwards burrowing lifestyle and termite-heavy diet, *Myobatrachus* lack many of the characteristics we typically associate with frogs. Their beady black eyes are set in small heads and, alongside their sister taxon *Arenophryne,* they crawl—not jump—across the ground on short limbs that are incapable of hopping (Vidal-Garcia et al. 2014).

## Conclusion

Australian frogs offer important insights into colonization, persistence, and diversification of a major continental group through deep time. The varied species richness, timing of diversification, and ecomorphological diversity among replicate radiations provides evidence of the processes dictating the accumulation of biodiversity. Beyond the temperate and tropical forests of the east and north coast, the Australian continent is an open country of habitat scarcely welcoming to frogs. Despite this, anurans have a long history in Australia and have diversified into an amazing array of forms, colors, and lifestyles. This success is potentially the result of the stratified temporal arrival of the three main frog clades and possibly exaggerated by their ecological differences. Our phylogenetic framework provides a foundation for further examining how temporal changes to climate, habitat, and niche space have influenced the diversification of one of Australia’s richest and most unique vertebrate faunas.

## Data Accessibility

Sequence alignments, analysis control files, and phylogenetic trees can be downloaded from Dryad (doi:10.5061/dryad.zpc866tcj) and GitHub (https://github.com/IanGBrennan/Crown_Frogs).

## Acknowledgments

Thank you to colleagues and staff at Australian museums and more generally across Australia for generously donating tissues and locality information for many frogs. We also thank the technical staff at our institutions for their support and hard work generating the genetic data presented here. The contributions of our many communities have made this work possible. JSK, CJH, and SCD thank the Australian Research Council for ongoing support. We appreciate comments from Isabel Sanmartín, Rayna Bell, and two anonymous reviewers that helped to improve a previous version of this manuscript.

## Supplementary Materials and Methods

Data available from the Dryad Digital Repository: http://dx.doi.org/10.5061/dryad.[NNNN] and from the GitHub repository: https://github.com/IanGBrennan/Crown_Frogs

### Developing Figure 1

Figure 1 aims to provide background on the richness and spatial distribution of the focal frog clades, alongside evolutionary context for the accumulation of vertebrate biodiversity on the Australian continent. Neither the top or bottom visualizations are intended to provide an explanation of the *processes* dictating Australian vertebrate diversity. Instead they are visualizations of the *patterns* of contemporary Australian vertebrate diversity.

We downloaded Australian annual rainfall data from NASA using the R package *nasapower*, and combined this with species occurrence records downloaded from the Atlas of Living Australia. Annual rainfall is an easily interpretable measure of water availability in an environment, and as such provides a reflection of habitat suitability for frogs. However, we acknowledge that composite environmental variables such as actual evapotranspiration (AET) may be a better predictor of contemporary frog richness patterns (Powney et al., 2010; Coops et al., 2018).

To plot the relationship between clade age and richness of Australian terrestrial vertebrates we collected data from all available non-nested (each clade is only represented once) clades from the literature. Data are compiled in the supplement *Comparative_Radiations.csv* and can be plotted using the script *Comparative_Radiations.R*. We also incorporated information where available about the biogeographic origin of each group to visualize the contrast between young clades from Asia and old Gondwanan groups. The included regression helps to visualize an interesting *pattern* in the data: species richness increases with clade age. However, we do not present this as an evolutionary explanation for varied richness among Australian terrestrial vertebrate groups.

### Sequence Identity

To confirm sequence identity we downloaded a fasta file of *Xenopus* genes from Ensembl (UCB_Xtro_10.0) and used *metablastr* to do a reciprocal blast against the Anchored Hybrid Enrichment loci. Of the 450 loci, 390 matched to *Xenopus* exons, and the remainder to intronic and flanking sequences (see *RBH_AHE_Xenopus.csv* in Supplementary Material for list). Downstream divergence time analysis relied on partitioning loci by codon position and so only exonic targets were retained for this exercise. AHE exons are listed under the column *query_id* and *Xenopus* matches under *subject_id* with gene name indicated by *subject_id_name*.

### Phylogenetics

Phylogeny reconstruction in the era of phylogenomics has simultaneously resolved many longstanding systematic questions and instigated new ones. The search for the most accurate species tree has reignited debates about concatenation versus coalescent methods and their pros and cons. Here we address two common issues resulting in phylogenetic error: incomplete lineage sorting (ILS) and rate variation among lineages and sites (heterotachy). Identifying and modelling heterotachy generally requires long alignments to accurately model rate variation, so most methods rely on concatenated sequence alignments. Because of the ancient age of our focal group and sparse sampling among major groups we risk biases due to heterotachy. To estimate a species tree from our concatenated alignment we used the General Heterogeneous evolution On a Single Topology (GHOST) method. GHOST is implemented in IQTREE and requires a user specified number of mixture (rate) classes and model. We separately fit unlinked GTR models with 2—5 mixture classes (e.g.: *-m GTR*H4*). AIC comparison identified the 4-class model as preferred (*H*2* AICc = 13754122; *H*3* AICc = 13604562; *H*4* AICc = 13500200; *H*5* AICc = 13523685).

Concatenation methods are however expected to perform poorly when the true branching pattern includes nested rapid divergence events. In this case high rates of ILS may bias phylogenetic signal, trapping concatenation in the anomaly zone. To counter this we estimated a species tree using ASTRAL with IQTREE genetrees as input.

### Biogeography

To assess the biogeographic history of Australian frogs we combined our phylogenetic hypothesis with known fossil information and reconstructed ancestral ranges in *BioGeoBEARS* (Matzke 2014). We started by dividing the geographic distribution of our sampled taxa into eight discrete areas that (1) summarize the general biogeographic history of frogs, (2) are relevant to our sampling and questions, and (3) make sense on a geological timescale with reference to plate tectonics over the last 220 million years. These areas correspond to Africa, Asia (excluding the Indian subcontinent), Australo-Papua, Europe, Madagascar, North America, South America, and Antarctica. For single tips that represent a genus or subfamily we coded their geographic range accordingly, however this never resulted in an overrepresentation of areas that might inflate dispersal estimates. Our primary objective was to identify the ancestral distributions of each Australian frog clade to provide an estimate of their origins.

While Antarctica seems a strange inclusion in our discrete bioregions owing to its current climate and lack of frogs, a recent discovery has identified the continent’s first anuran (Mörs et al. 2020). This information is vital to our understanding of the connectivity of the Gondwanan supercontinent as well as the biogeographic history of Australian frogs. To incorporate this sample we added a tip to our tree with an appropriate estimated age following Mörs et al. (2020). Due to our limited sampling of extant Calyptocephalellidae however, the addition of this taxon dramatically imbalances range reconstruction. To correct for this and account for the ancient known history of calyptocephalellids in South America (Moura et al. 2021; Nicoli et al. 2022) we included two additional South American fossil taxa, one younger—*Calyptocephalella canqueli* (following Muzzopappa & Báez 2009) and one older—*Calyptocephalella satan* (following Nicoli et al. 2022). Note, here we consider *C.satan* as interchangeable with the similarly aged *Baurubatrachus pricei* (following Báez & Gómez 2018), being representative of a broader extinct South American calyptocephalellid diversity (Nicoli et al. 2022). While the taxonomy and phylogenetic relationships of extant (*Calyptocephalella gayi*, *Telmatobufo spp.*) and extinct (*C. canqueli, C. satan, et al.*) calyptocephalellids is unresolved, we believe this sampling strategy is an appropriate solution for the question at hand.

In addition to the origins of Australian frogs we were interested in identifying how pelodryadids arrived in Australia. Specifically we aimed to test if they arrived via dispersal through Antarctica or overwater dispersal from South America. To test these hypotheses we added an ancestor (*Pelodryadidae_Ancestor*) to our tree along the stem leading to the Pelodryadidae. *BioGeoBEARS* accommodates sampled ancestors as “hooks”, which are represented by a non-zero terminal edge length shorter than an arbitrary threshold (here: 0.000001 million years). This allowed us to force the ancestral pelodryadid to either have had a range in Antarctica (Hypothesis 1; H1; South America→Antarctica→Australia), or have remained in South America prior to an overwater dispersal event (Hypothesis 2; H2; South America→Australia).

The biogeographic history of frogs has played out on a very long timescale (>200 million years) and across continents that have moved dramatically relative to one another. To capture the complex interplay of plate tectonics and biogeography we incorporated several elements that might make this scenario more realistic. We first divided the anuran tree into six equal slices of 30 million years (0—30, 30—60, … 150—180) and one slice of 40 million years (180—220). At the upper bound of each time slice (30, 60 … 180, 220) we then reconstructed continental positions in GPlates following Landis (2017) and extracted pairwise distances (in km) among areas from the closest points of two areas, using the measuring tool in GPlates. Additionally, we characterized regions as (a) in contact with one another, (b) separated by ocean, or (c) separated by another landmass. We used the area distances through time to construct distance matrices following Van Dam & Matzke (2016), and the area adjacency information to construct dispersal matrices.

Constructing these time-specific matrices allowed us to compare a set of scenarios that include the traditional DEC model (Dispersal Extinction Cladogenesis), DEC+j which allows jumps in range expansion (range discontinuity), DEC+x which estimates a parameter *x* corresponding to a correction for dispersal probability as a function of distance between areas (dispersal * relative_distance^*x*), DEC+j+x which allows jumps and corrects for distance among areas, DEC+x+w which estimates *x* (correcting for distance) in addition to a parameter *w* which can be interpreted as correcting for different levels of area adjacency (dispersal * dispersal_multiplier^*w*), and finally DEC+j+x+w which can be interpreted as allowing for jumps in range expansion (*j*) while correcting for geographic distance between areas (*x*) and types of adjacency/separation (*w*). Ultimately the most complex model (DEC+j+x+w) is an attempt to account for differences in the geographic distance between areas (*x*) as well as what separates them (*w*), through time, while allowing taxa to make rapid dispersal events (*j*). Estimating *w* unfortunately necessitates the manual input of dispersal multipliers which scale dispersal probability, however these are ultimately corrected by estimating their relationship via *w*. We established conservative manual dispersal multipliers for adjacent areas (1), areas split by another contiguous landmass (0.5), and areas split by ocean (0.25). Finally, we fit all six models to both the H1 and H2 datasets. We compared models by calculating AIC values, delta AIC against the best fit, and AIC weights as the relative contribution to the pool of models.

**Table S1.**
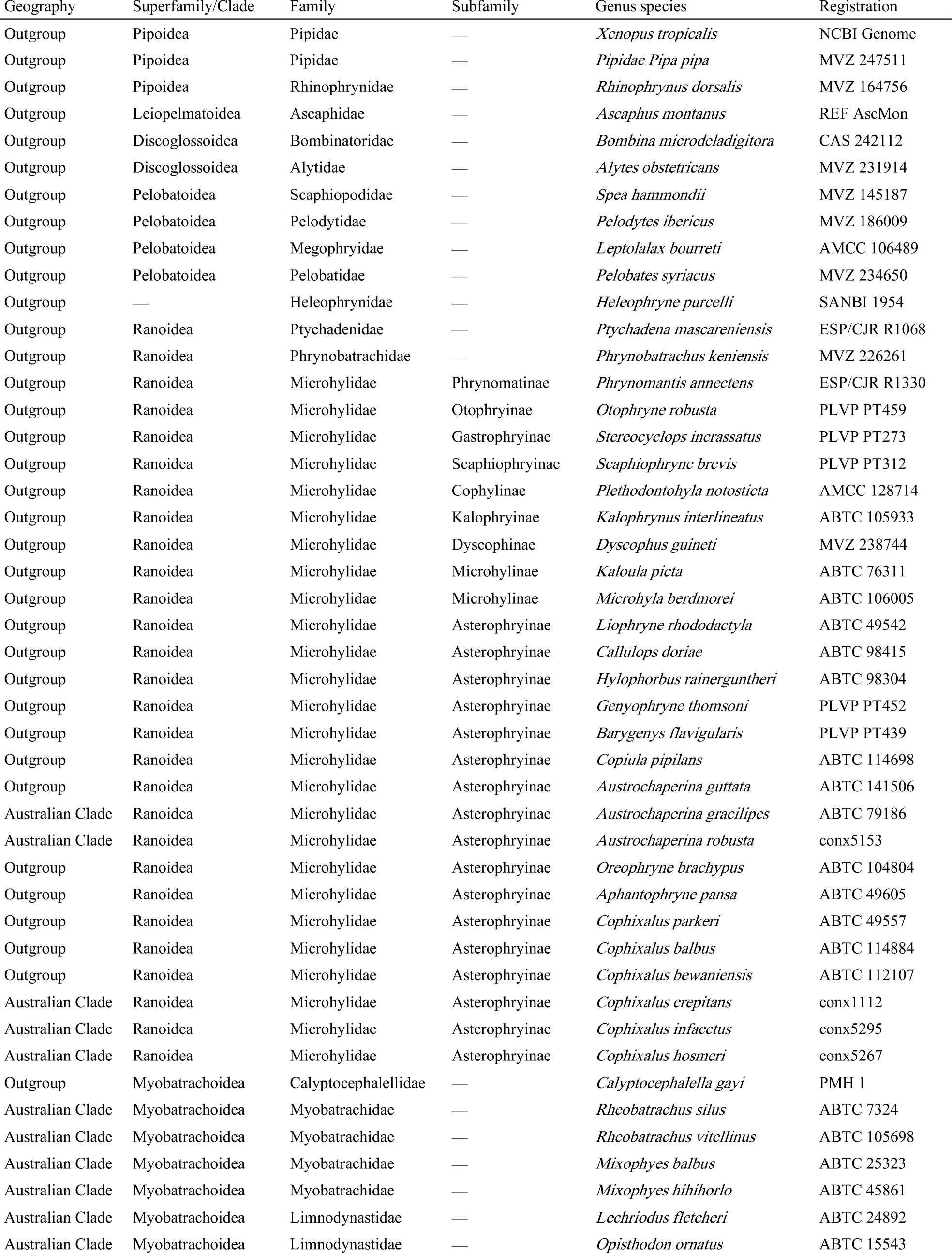
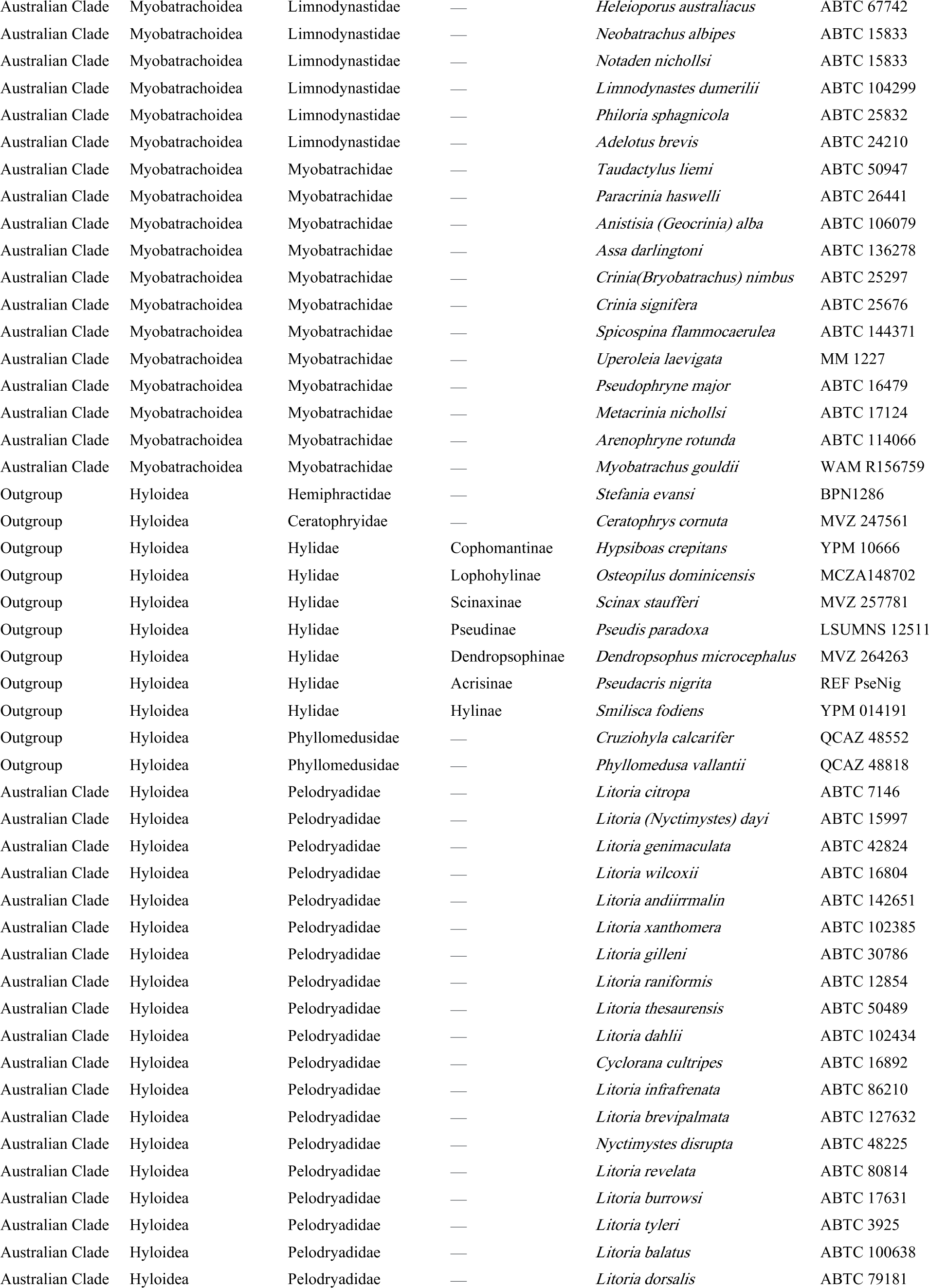
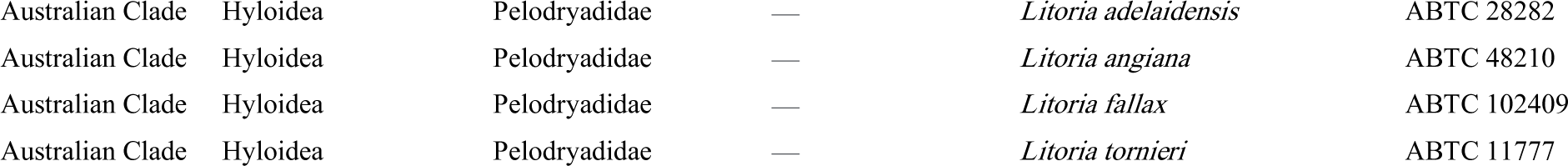
Taxon sampling for this project.

**Table S2.**
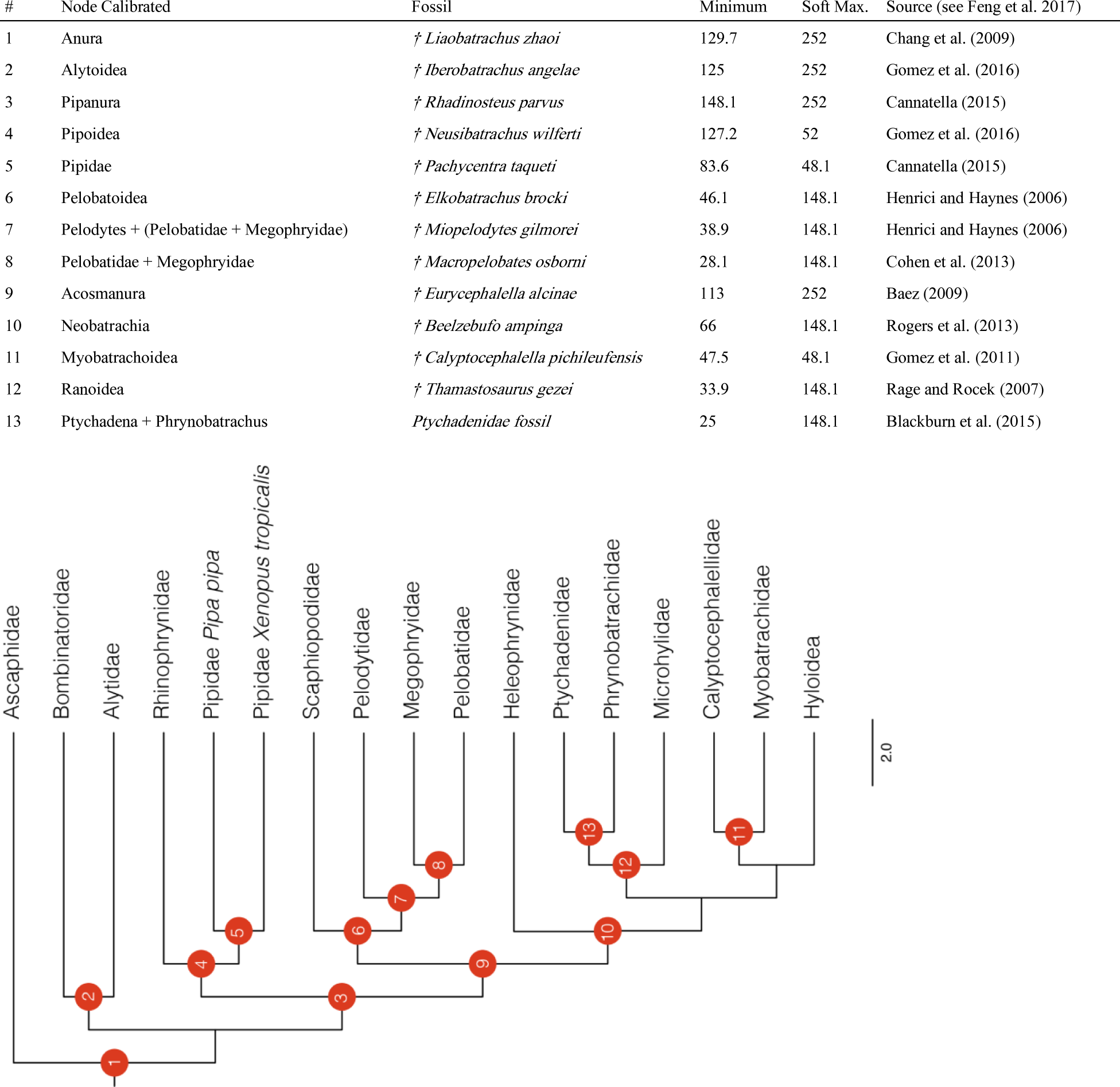
Fossil calibrations implemented in MCMCtree analysis of frog divergence dates. Node number (#) corresponds to nodes in supplementary figure below.

**Table S3.**
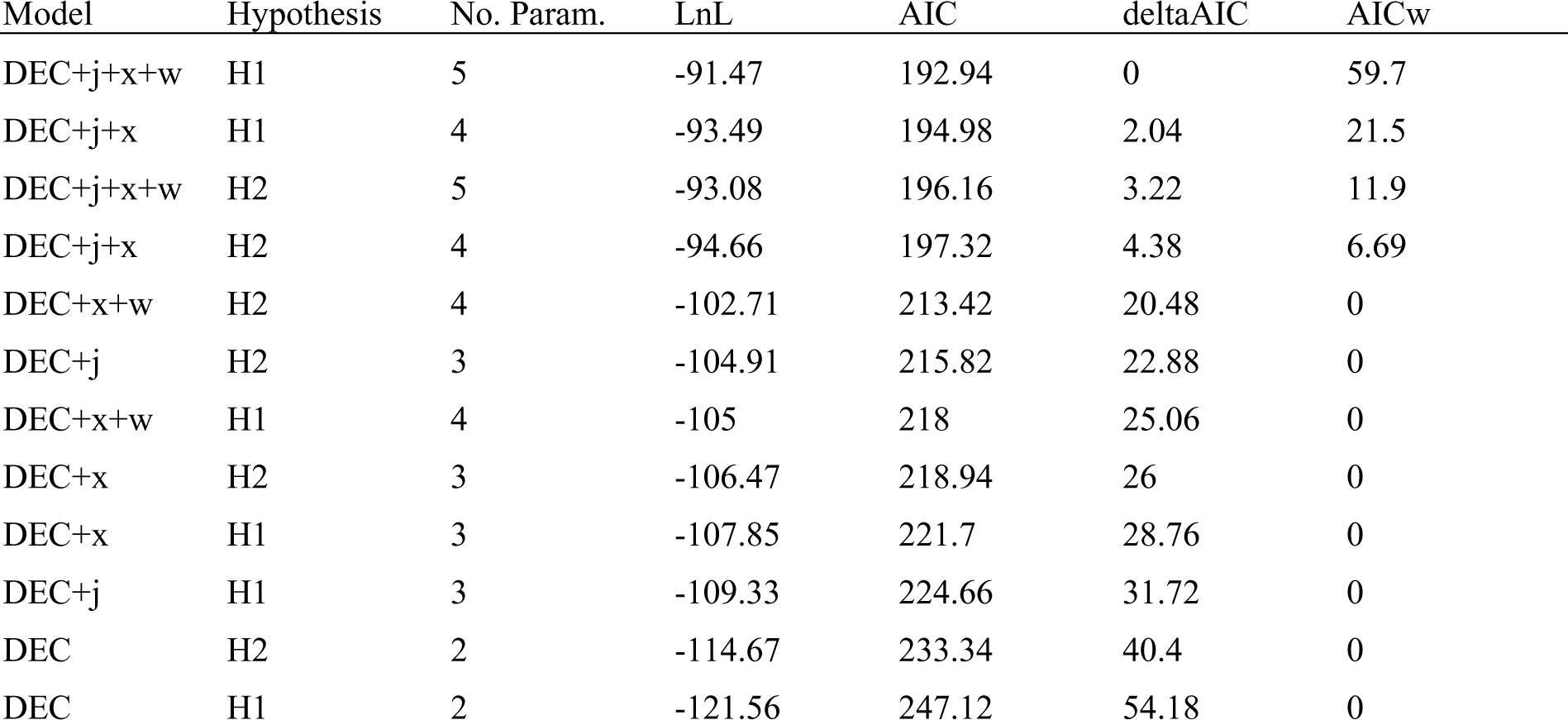
Results of biogeographic ancestral range reconstruction in *BioGeoBEARS*. Hypothesis *H1* refers to the dispersal of pelodryadid frogs from South America through Antarctica to Australia, whereas *H2* refers to the over water dispersal of pelodryadid frogs from South America directly to Australia. Models are sorted according to deltaAIC scores, indicating the preferred model at the top.

| Model | Hypothesis | No. Param. | LnL | AIC | deltaAIC | AICw |
| --- | --- | --- | --- | --- | --- | --- |
| DEC+j+x+w | H1 | 5 | -91.47 | 192.94 | 0 | 59.7 |
| DEC+j+x | H1 | 4 | -93.49 | 194.98 | 2.04 | 21.5 |
| DEC+j+x+w | H2 | 5 | -93.08 | 196.16 | 3.22 | 11.9 |
| DEC+j+x | H2 | 4 | -94.66 | 197.32 | 4.38 | 6.69 |
| DEC+x+w | H2 | 4 | -102.71 | 213.42 | 20.48 | 0 |
| DEC+j | H2 | 3 | -104.91 | 215.82 | 22.88 | 0 |
| DEC+x+w | H1 | 4 | -105 | 218 | 25.06 | 0 |
| DEC+x | H2 | 3 | -106.47 | 218.94 | 26 | 0 |
| DEC+x | H1 | 3 | -107.85 | 221.7 | 28.76 | 0 |
| DEC+j | H1 | 3 | -109.33 | 224.66 | 31.72 | 0 |
| DEC | H2 | 2 | -114.67 | 233.34 | 40.4 | 0 |
| DEC | H1 | 2 | -121.56 | 247.12 | 54.18 | 0 |

**Figure S1.**
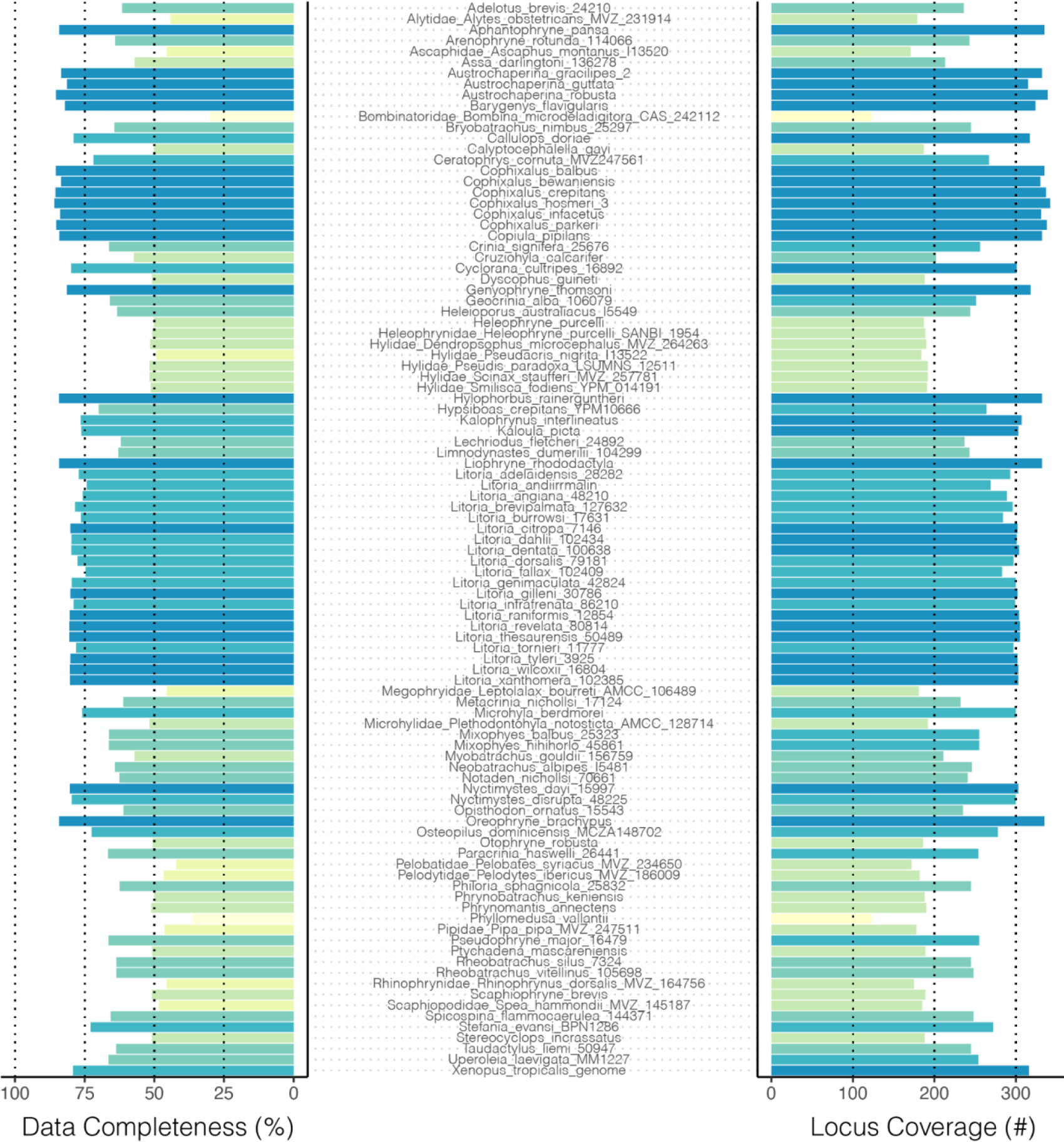
Data completeness across all samples. Left histogram shows data completeness as percent of bases in total alignment (concatenated alignment length 523,036 bp) exclusive of gaps (-) and missing bases (N). Right histogram shows data completeness as the absolute number of loci included per sample, as a representation of the number of gene trees per sample.

**Figure S2.**
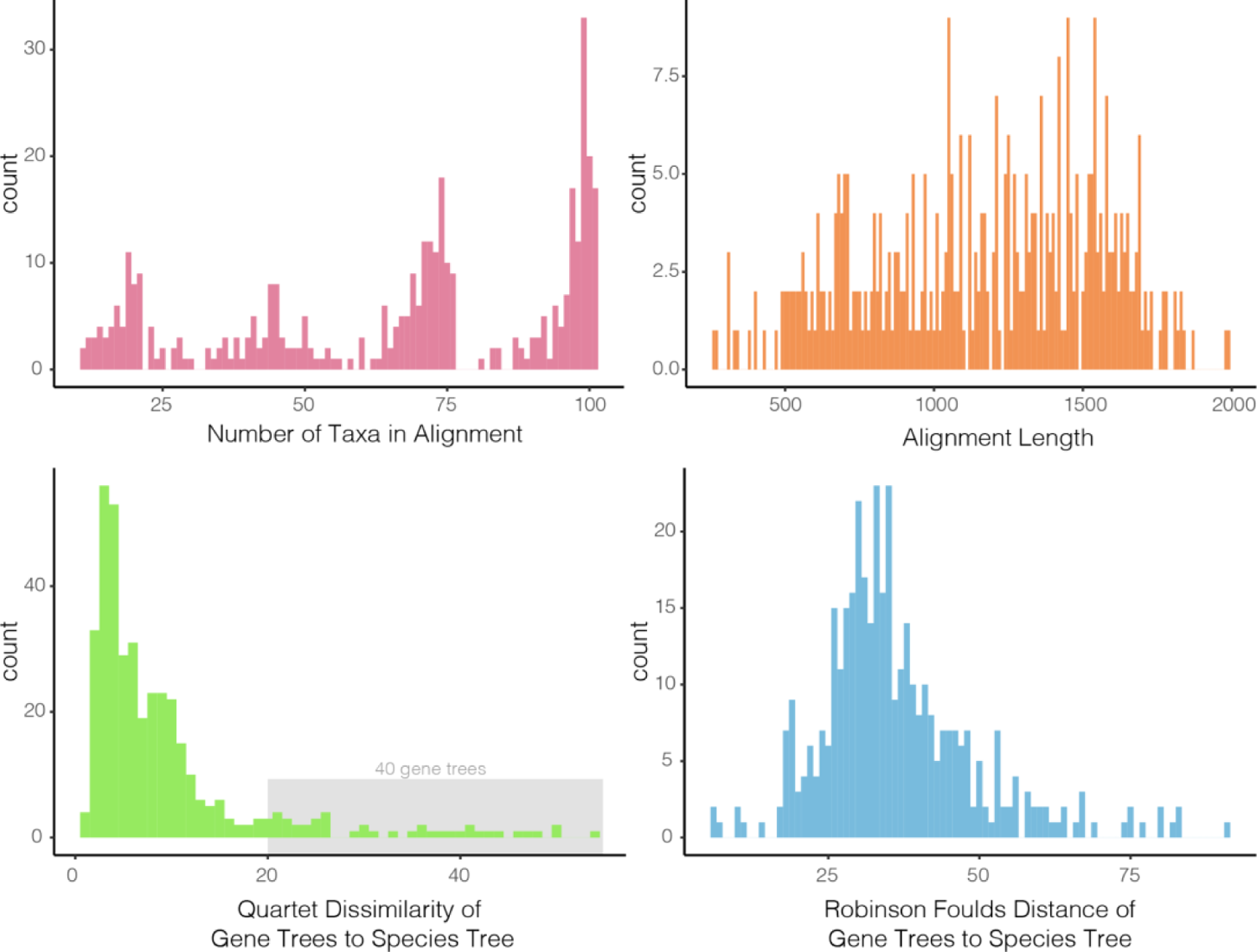
Basic summary statistics of the 450 locus alignments and gene trees. Top row shows histograms of the number of taxa in (max=101, min=11) and length of each alignment. Bottom row shows gene tree--species tree distances as quartet dissimilarity scores and Robinson Foulds distances, two different measures of topological similarity. Both quartet dissimilarity and RF scores are estimated by first subsetting the species tree to match gene tree sampling.

**Figure S3.**
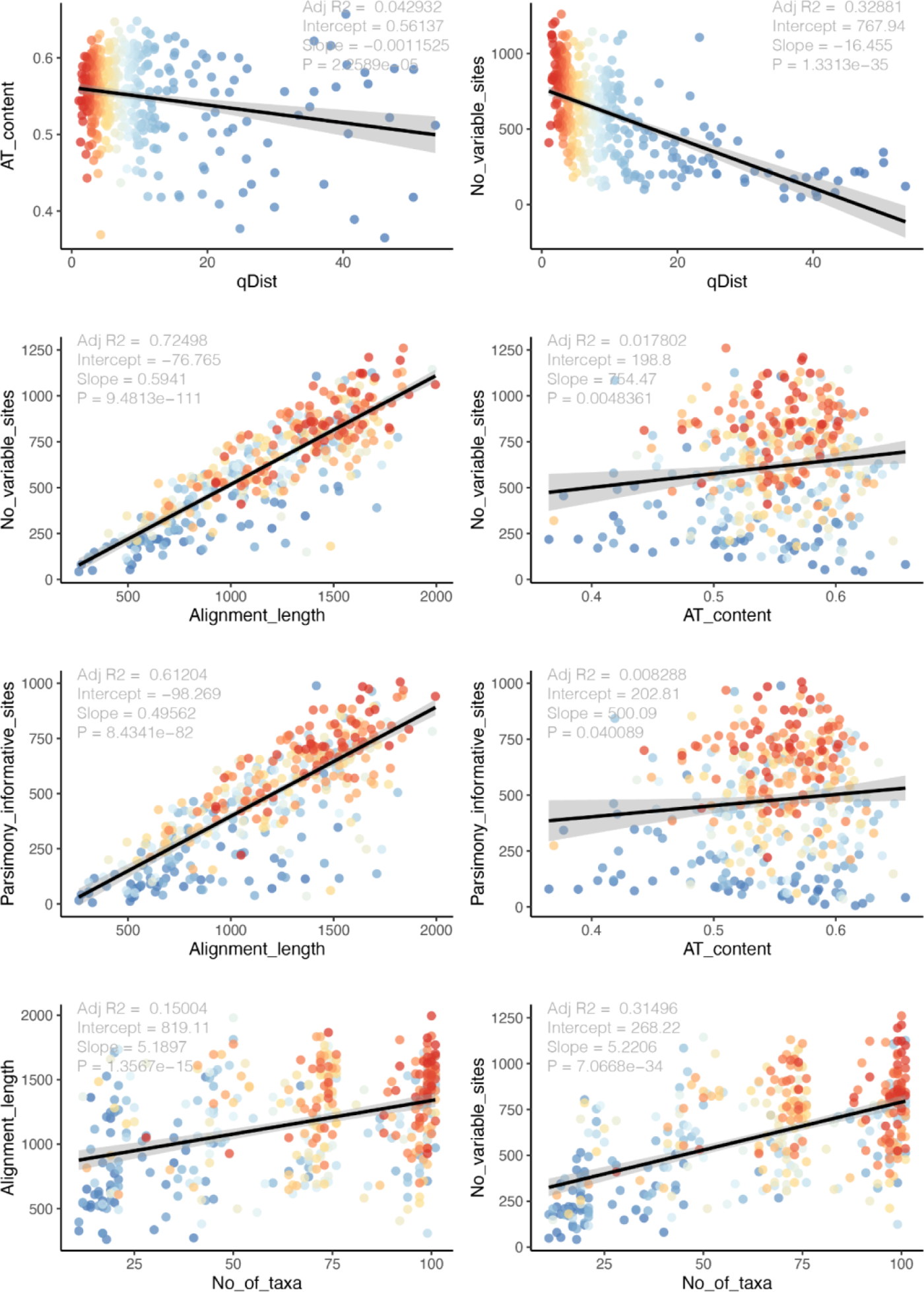
Detailed summary statistics of the 450 locus alignments and gene trees. Top row compares AT content and number of variable sites against quartet distance between each gene tree and the species tree (a measure of topological similarity). The second and third rows compare measures of locus informativeness (number of variable sites, number of parsimony informative sites) against alignment length and AT content. The bottom row shows alignment length and number of variable sites as a function of the number of taxa in the alignment. In all plots points (representing trees or alignments) are colored according to the quartet distance from the species tree.

**Figure S4.**
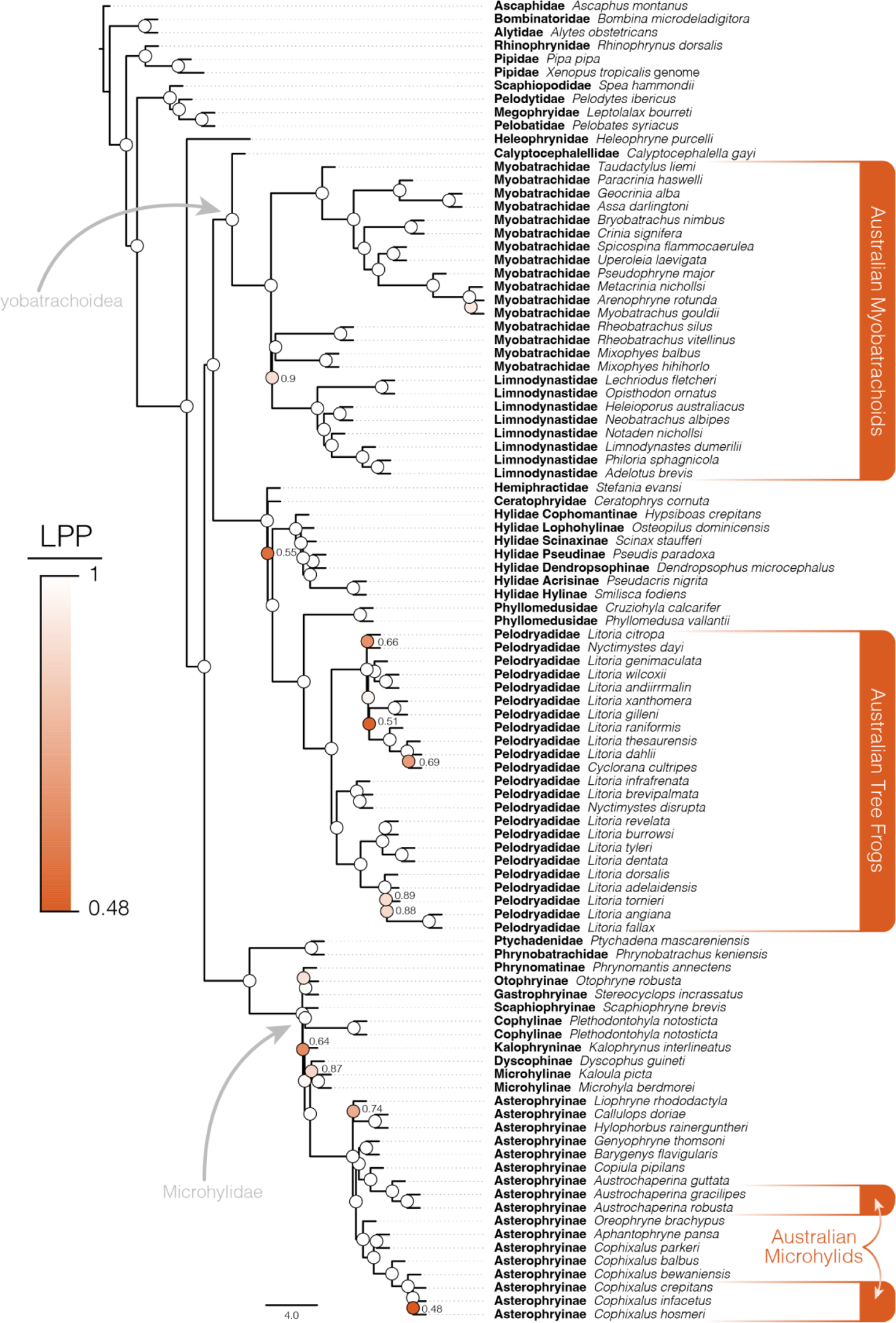
Species tree of Australian frogs and appropriate outgroup taxa estimated using ASTRAL with locus trees estimated by IQTREE as input. Phylogenetic resolution among major frog groups and within Australian frog clades is consistently high. Ultrafast bootstratp support values (Hoang et al. 2018) are shown at nodes and colored according to local posterior probabilities (LPP), values >0.9 are considered strongly supported and not indicated at nodes (white circles).

**Figure S5.**
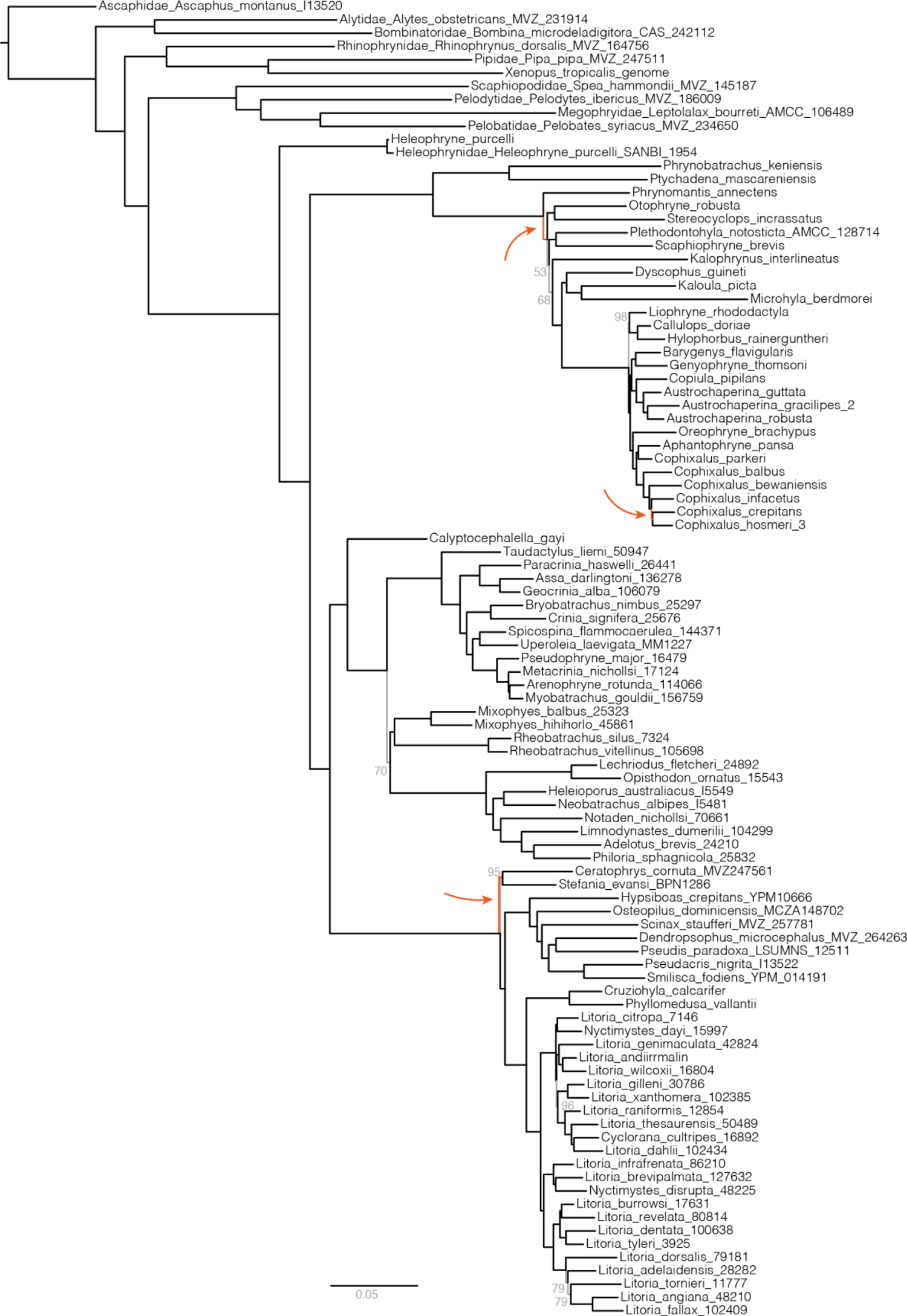
Species tree of Australian frogs and appropriate outgroup taxa estimated from the concatenated sequence alignment under the GHOST model implemented in IQTREE. Phylogenetic resolution among major frog groups and within Australian frog clades is consistently high. Only ultrafast bootstrap support values less than 100 are noted, here by grey branches and text (Hoang et al. 2018). This topology is highly consistent with the phylogeny estimated using ASTRAL (Fig.2, S4), however three differences are highlighted by orange branches and arrows indicating their location. Branch lengths are weighted averages over four heterotachy classes.

**Figure S6.**
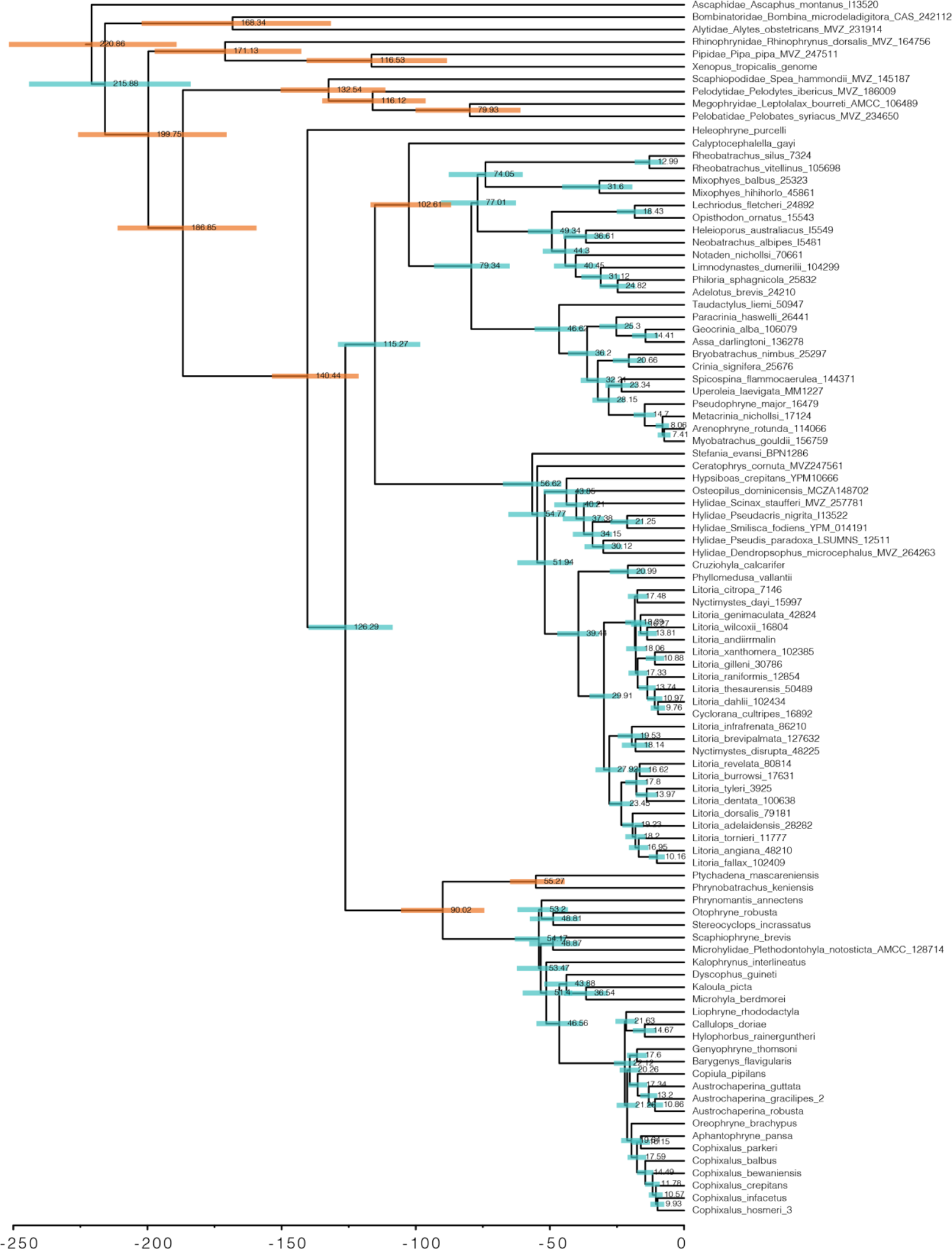
Species tree of Australian and outgroup frogs estimated with ASTRAL from IQTREE genetrees and time-calibrated with MCMCtree. Shaded bars at nodes indicate 95% confidence estimates on ages and numbers indicate mean age estimates. Orange shaded bars indicate nodes which were calibrated with from fossil evidence (see Table S2).

**Figure S7.**
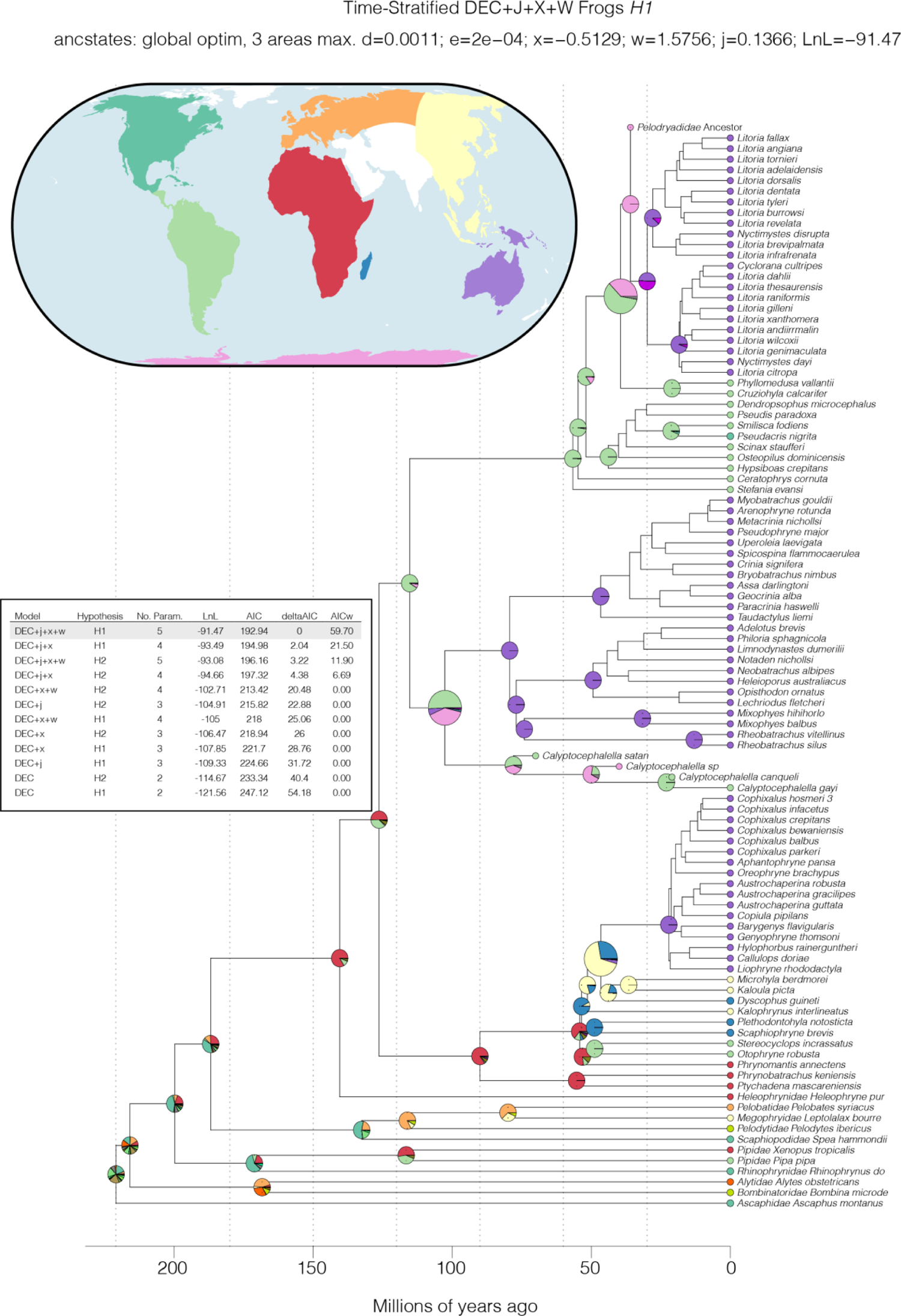
Biogeographic history of frogs with a focus on the range reconstruction of Australian clades. Inset table shows the 12 models fit to the data (6 models across two ‘datasets’), ordered by deltaAIC. Ancestral range estimates under the preferred model DEC+*j*+*x*+*w H1* are shown at right as pie charts on the phylogenomic tree with several fossil taxa added. Pie chart for the most recent common ancestor of each Australian clade is enlarged to enhance visualization. The eight bioregions are shown in the inset map and colors correspond to the tip state of taxa on the tree. Additional colors in the pie charts correspond to combinations of areas, but are not discussed further.

